# Volitional generation of reproducible energy-efficient temporal patterns

**DOI:** 10.1101/2022.04.27.489830

**Authors:** Yuxiao Ning, Guihua Wan, Tengjun Liu, Shaomin Zhang

## Abstract

One of the extraordinary characteristics of the biological brain is its low energy expense to implement a variety of biological functions and intelligence compared to the modern artificial intelligence (AI). Spike-based energy-efficient temporal codes have long been suggested as the contributor for the brain to run with a low energy expense. Despite this code having been largely reported in the sensory cortex, whether this code can be implemented in other brain areas to serve broader functions and how the brain learns to generate it have remained unaddressed. In this study, we designed a novel brain-machine interface (BMI) paradigm, by learning which two macaques could volitionally generate reproducible energy-efficient temporal patterns in the primary motor cortex (M1). Moreover, most neurons that were not directly assigned for controlling the BMI did not boost their excitability, demonstrating an overall energy-efficiency manner in performing the task. Over the course of learning, we found that the firing rates and temporal precision of selected neurons co-evolved to generate the energy-efficient temporal patterns, suggesting a cohesive rather than dissociable processing underlie the refinement of energy-efficient temporal patterns.

## Introduction

Although the human brain actively sapped 20% of body energy, it ultimately consumes only 20 watts of power (Raichle and Gusnard, 2002; Sokoloff, 1960). Moreover, one newest study revealed that power expended for computation takes less than one percent of the total budget (Levy and Calvert, 2021). Compared to the amount of energy consumed by modern artificial intelligence (AI) implemented on silicon-based hardware, the brain consumes extraordinarily little to implement biological intelligence. For decades, scientists and engineers have attempted to draw insights from neural coding schemes to build energy-efficient intelligent systems. Temporal coding, by which information is communicated and processed through the temporal coordination of spikes, utilizes the discrete nature of spikes. It has been proven that the temporal code mathematically carries more information than the rate code (Rieke et al., 1999; Denève and Machens, 2016). Therefore, temporal coding has grabbed attention and has recently been employed in AI practices (Bohte et al., 2002; Mostafa, 2017; Comsa et al., 2020). However, the implementation of temporal codes does not necessarily lead to sparsity or a low energy budget globally in biological brains. Riehle et al. (1997) has found that precise synchronization of spikes could occur both in absent and in the presence of firing rate elevation. In addition, the study by Ning et al. (2022) indicated that the excitatory gain of the population propelled the neural network to a regime that facilitates synchronization and the generation of precise temporal patterns. Therefore, a deeper understanding of how energy-efficient temporal patterns are generated and the neural difference underlying the generation of efficient and non-efficient patterns is much needed. This is the keystone to drawing functional implications in the realm of neuroscience and guiding the design of next-generation artificial intelligence and neuromorphic architectures.

Despite many reports that have observed energy-efficient temporal patterns in the sensory system and related them with different stimuli or behaviors (Panzeri et al., 2001; Arabzadeh et al., 2006), our understanding remains incomplete. First, the representational capability of energy-efficient temporal patterns is well acknowledged due to studies conducted in the sensory system, but whether the energy-efficient temporal patterns have the computational capability is far from being explored and understood. The computational capability of energy-efficient temporal patterns is also of great interest in AI and neuromorphic computing (Lansdell and Kording, 2019). However, if we want to draw biological inspiration on this computational capability, we have to extend the study of energy-efficient temporal patterns to other brain areas where more diverse and complex computations operate, not being limited to perception in sensory areas. Unfortunately, studying the implementation and computation of efficient temporal patterns in brain areas other than the sensory cortex can be hampered by the representational view and the derived behavioral paradigms. For example, it is difficult to assign semantic meaning to neural patterns in high-level cortices because the context there would be more abstract and entangled (Asaad et al., 1998; Rigotti et al., 2013). Besides, recent studies indicated that neural activity in motor cortices can be a reflection of its own dynamics rather than representing detailed kinematics (Churchland et al., 2010, 2012; Shenoy et al., 2013). Therefore a new paradigm for studying the implementation of efficient temporal patterns on various cortices is needed. Second, it is unclear how the energy-efficient temporal patterns, where changes from single spikes matter, are learned and evolved. Understanding this underlying learning is yet critical for developing training algorithms of spiking neural networks (SNN), since some of the well-developed algorithms for training traditional neural networks, such as gradient descent, cannot be directly transferred to SNN for the indifferentiability of discrete events. However, it is challenging to study the learning and evolution of energy-efficient temporal patterns in biological brains. Without knowing the causality from temporal patterns to behaviors, one can hardly infer the functional relevance of changes on single spikes.

Aside from its resounding success in restoring motor (Nicolelis, 2001; Carmena et al., 2003; Hochberg et al., 2006), speech (Moses et al., 2021), and psychological functions (Shanechi, 2019), brain-machine interfaces (BMIs) has recently energized neuroscience by offering causality for certain neural patterns to behavioral outcomes. Scientists can define the mappings between neural activity and outcomes, which was termed as “decoders” (Sadtler et al., 2014; Hennig et al., 2018; Golub et al., 2018; Clancy and Mrsic-Flogel, 2021). In the study of Ning et al. (2022), the authors showed that when the decoders operate on the temporal domain rather than the commonly used descriptors like firing rates, to control outcomes, the elicited patterns with temporal precision can be causally related to behaviors. Besides, known mappings in such “temporal neuroprosthetics” allow them to track how the temporal patterns are learned and refined in brain circuits. However, they only covered how nonefficient temporal patterns were generated and learned, leaving energy-efficient temporal patterns largely unexplored. Therefore, in this study, we introduced a new BMI and studied the generation and learning of energy-efficient temporal patterns in motor cortex.

## Results

We devised a novel BMI paradigm to explore the plausibility of generating energy-efficient temporal patterns in M1. The principle for designing the paradigm was to encourage higher temporal precision while penalizing excessive firing. To this end, two key ingredients in a BMI system should be purposely designed: the “decoder” and the feedback approach. For the BMI “decoder”, we developed the normalized coincidence score (NCS) as the decoded variable (Methods). The numerator of the NCS, termed as the coincidence score, served the function of rewarding for not only the temporal precision but also the correct temporal order. It is a function of the relative timing between the trigger unit and the target unit, which took the form of a Spike-timing dependent plasticity (STDP) function. Next, the coincidence score was normalized by the geometric mean of spike counts from the two units to render the NCS, which is equivalent to penalizing excessive spikes. Unlike the widely used BMI “decoders” based on linear time-invariant mappings, ours based on nonlinear mapping might incur greater complexity. Thus, we adopted auditory and visual feedback simultaneously in the task to complement each other.

The duration of the learning course varied in two subjects (B11: 10 sessions, C05: 18 sessions). However, for analytical and visualizational convenience, we truncated C05’s course of learning into identical lengths with B11 by removing intermediate sessions (Session 6 to Session 13). Furthermore, we designated the first half of the sessions into the early group and the last half into the late group. We found that both subjects exhibited a certain improvement on this task. The normalized success rate increased over the course of learning (Fig. 2A, 2C), while the trial duration decreased (Fig. 2B, 2D). Besides, the distribution of NCS was flatter in the task block than that in the baseline block (Fig. 3A *left*). This disparity became even more prominent in the late phase of learning, indicating that the subjects mastered the modulation of NCS in the late block of the course (Fig. 3A *right*). Similar change was also observed when comparing the distribution of NCS from task block in the first session with that in the last session. (Fig. 3B) (B11: *var*_*S*1_ < *var*_*S*10_, *F* = 0.4525, p < 0.0001; C05: *var*_*S*1_ < *var*_*S*10_, *F* = 0.7304, p < 0.0001). Collectively, the novel task we designed was learnable, under which the subjects became proficient in modulating the non-linearly mapped NCS.

**Figure 1.**
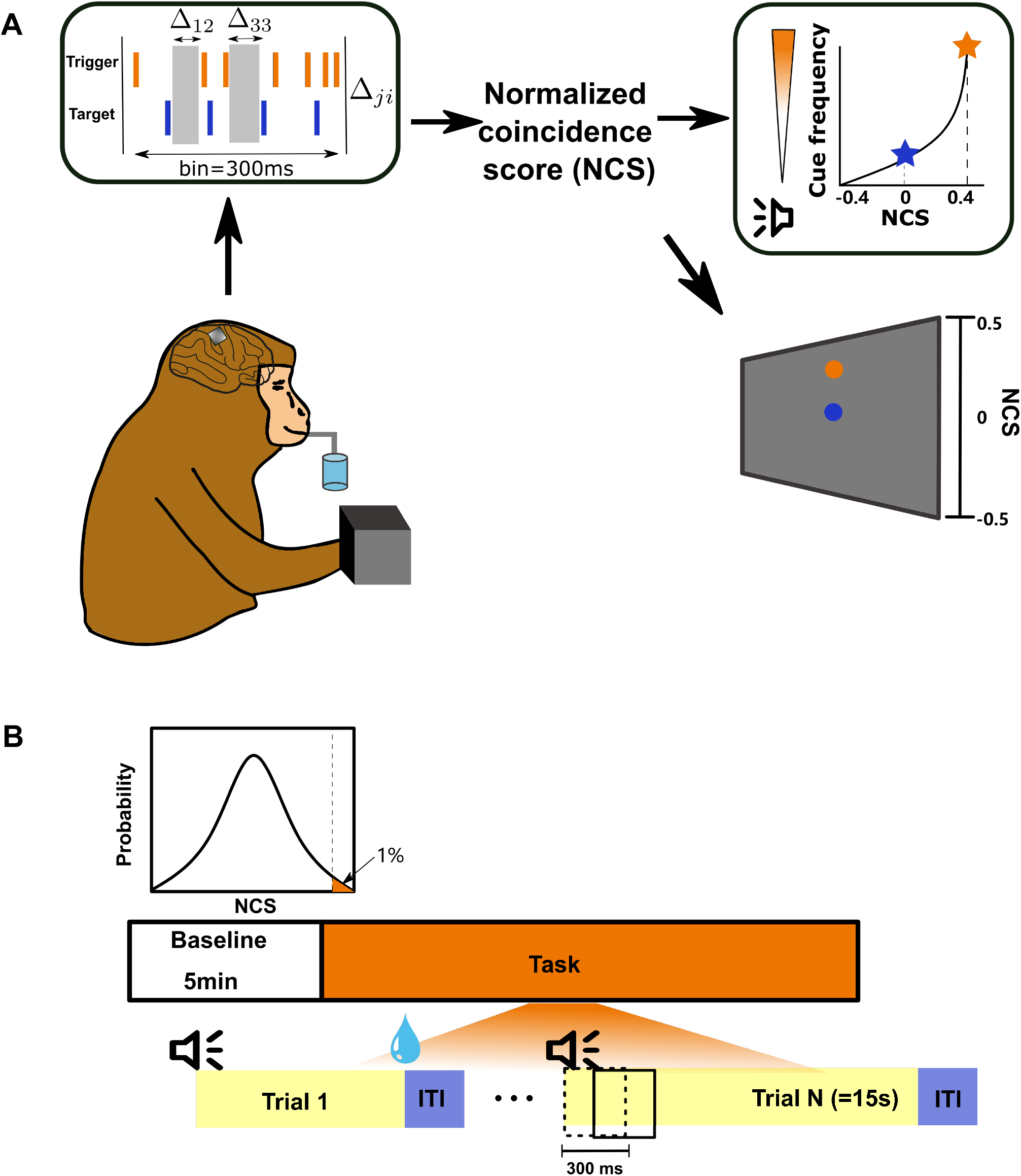
The BMI-based task paradigm. (**A**) The task was run in a closed-loop fashion. Neural activity from subject’s M1 was automatically read and extracted. All the leads and lags between spikes from trigger and target units were used to computer NCS, which was further fed back by mapping it to the frequency of an audio cursor and the position of an visual cursor. In this example trial, the NCS threshold for reward is 0.4. Subject’s hands were constrained through the training session. (**B**) Task structure of one typical session. The session was kick-started by a 5-minutes baseline block, after which the distribution of NCS could be estimated. The threshold for water reward was set on the basis of the NCS distribution estimated in the first session (inset). One trial could span to at most 15 s followed by a 4 s-long intertrial interval. The NCS was computed using neural activity in a 300 ms sliding window.

**Figure 2.**
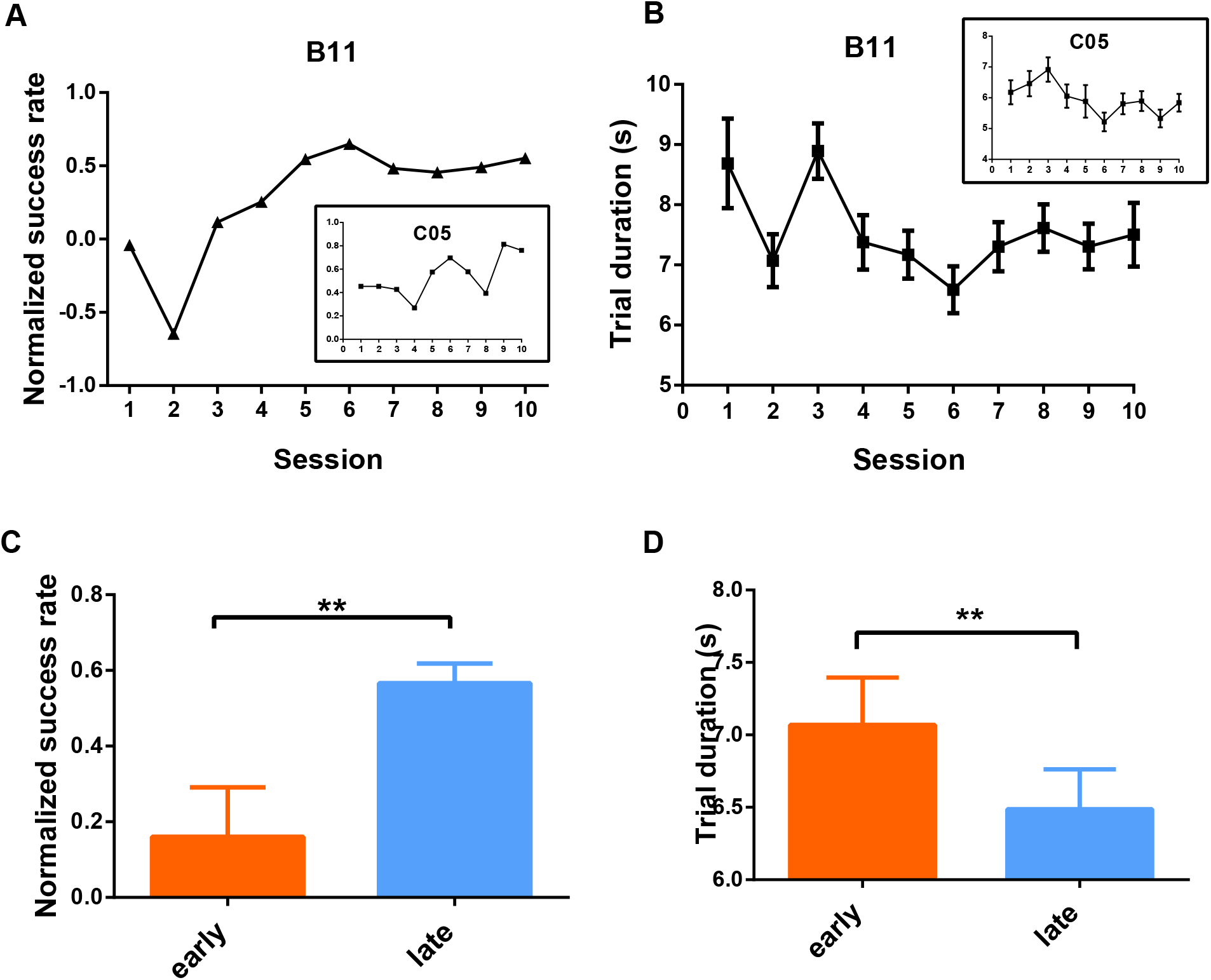
Proficiency increased in performing the BMI-based task. (**A**) The normalized success rate exhibited a rising trend over 10 sessions. (**B**) The trial duration reduced over 10 sessions. (**C**) Pronounced increase in the normalized success rate from the early block to the late block, pooling over two subjects. Mean ± SEM. Unpaired t-test with Welch’s correction, early < late, p = 0.008. (**D**) Significant shortening in trial duration from the early block to the late block, pooling over two subjects and all trials. Mean ± SEM. F(1,2335) = 10.41, p = 0.0013.

**Figure 3.**
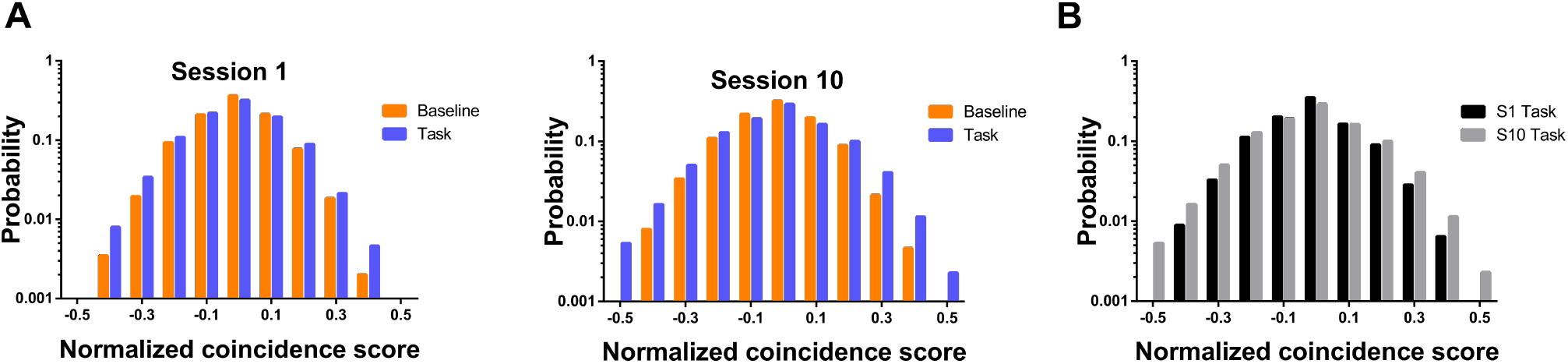
Modulation of NCS became stronger in late phase of learning. (**A**) C05’s NCS distribution exhibited modulation of NCS in the task block. The Y-axis was set as the logarithmic scale to better reveal the changes of NCS around the rewarding threshold. (**B**) C05’s NCS distribution in task block at the first session and the last session.

We next asked whether up-modulating NCS could evoke the precise temporal pattern in this task. To reveal the precise temporal patterns, we applied the cross-correlation histogram (CCH) to the 300 ms preceding reward delivery (denoted as “Pre-reward”). We first binned the CCH in 15 ms, consistent with the time constant of exponential decaying in the STDP function. The CCH peaked near zero exclusively on the positive side, suggesting a notable high probability of spike coincidence and a reliable temporal order (Fig. 4A). Furthermore, since the coincidence score was exponentially decayed with increased relative spike timing, subjects were expected to generate finer temporal patterns (≤ 5 ms) to improve the chances of obtaining rewards. By narrowing the bin width of the CCHs to 5 ms, we found that the target units indeed fired more compactly with the trigger units than what could be inferred from the coarser CCHs mentioned above (Fig. 4B).

**Figure 4.**
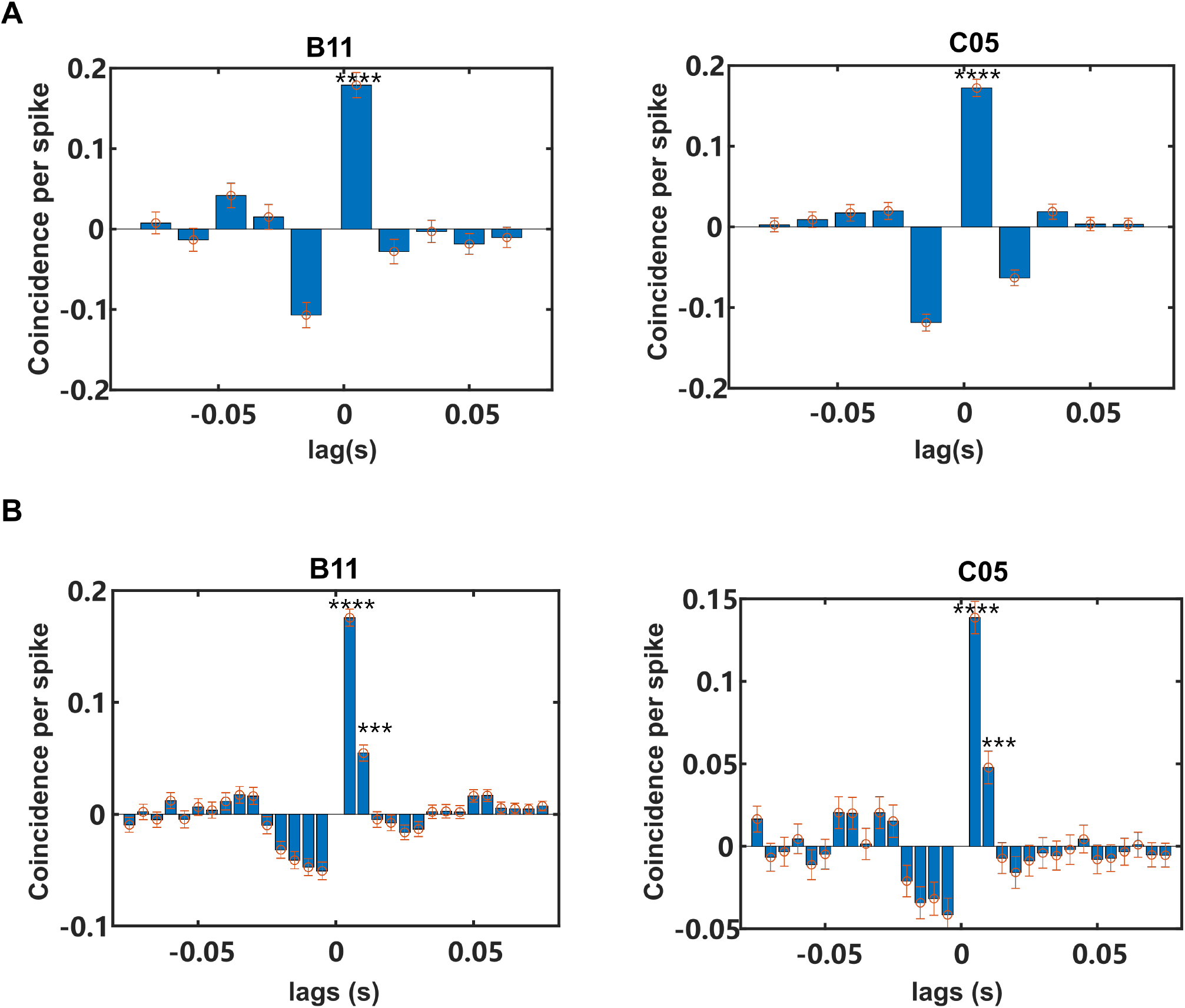
Temporal precision increased while learning the BMI-based task. (**A**) CCHs of two subjects in the last session showed pronounced coincidence between two units in time window spanning from 0 to +15 ms. Error bars denoted the SD rendered from resampling analysis on jittering. The asterisks labeled above the bars indicated a significant coincidence. ****: p<0.0001. (**B**) The CCHs of two subjects in the last session with higher resolution showed finer temporal granularity. Error bars denoted the SD rendered from resampling analysis on jittering. The asterisks labeled above the bars indicated significant coincidence. ***: p<0.001, ****: p<0.0001

Additionally, the main purpose of normalizing the coincidence score was to keep the occurrence of discharging events under a low level which served to cut the energy expenses. Thus, we analyzed how the firing rates of the selected pair of neurons changed across sessions. We found that over the course of learning, the change in firing rates of the pair was brought largely by the target units (Fig. 5A, test for the significantly non-zero slope of linear fitting, B11: trigger unit, p = 0.354; target unit, p = 0.004; C05: trigger unit, p = 0.705; target unit, p = 4e-4). On top of that, the variation of firing rates across sessions was identical between data from baselines and tasks for target units but not trigger units (Fig. 5A, Pearson test for correlation, B11: trigger unit, r = 0.388, p = 0.267; target unit, r = 0.923, p = 1e-4; C05: trigger unit, r = −0.263, p = 0.462; target unit, r = 0.960, p = 1e-4). We thus investigated how the firing rates of the direct units changed compared to the baseline level. Modulation index (MI) was commonly used to measure the relative change of firing rates under distinct conditions normalized by the average firing level (Movshon et al., 1978; Ganguly et al., 2011). Since down-modulating of firing rates was more of a concern in our case, we modified the MI into the directional modulation index (d-MI, Methods). Significant modulation to reduce discharging events would result in a positive d-MI, while greater increase of firing rates relative to the baseline level would lead to a more negative d-MI. Fig. 5B showed the MI of the target units enhanced at the last session, suggesting a stronger reduction in spike counts. Therefore, together with the temporal precision revealed in Fig.4, we demonstrated that energy-efficient temporal patterns between two arbitrarily assigned neurons in M1 could be generated by learning the novel BMI paradigm.

**Figure 5.**
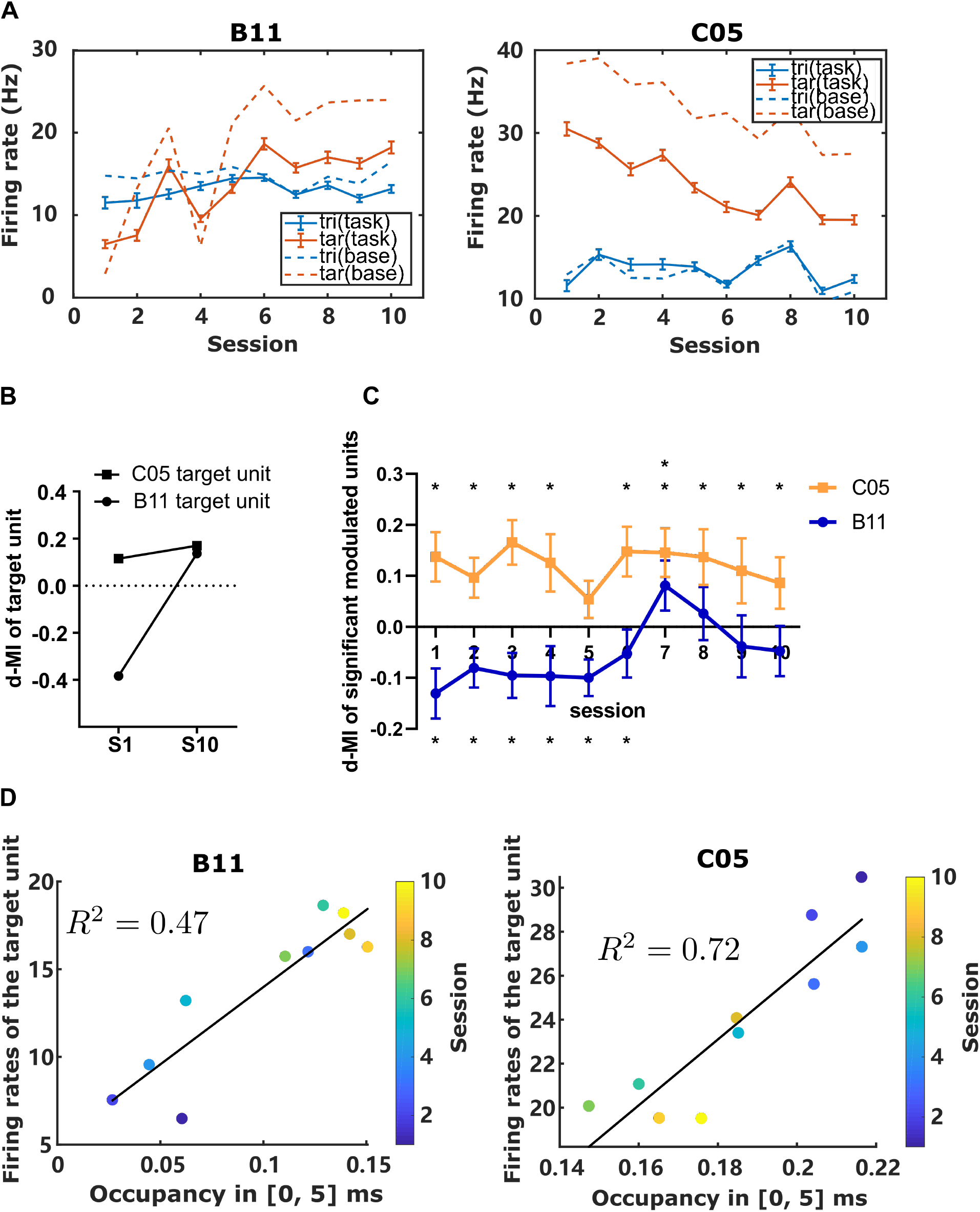
Efficient firing of both the direct neurons and indirect neurons. (**A**) The evolution of firing rates of direct neurons in learning. Color was coded for different neurons, while line type coded for different blocks in one session. (**B**) The d-MI of target unit significantly increased. (**C**) Average d-MI after pooling over all significantly modulated indirect neurons across sessions (mean ± SEM). The asterisks labeled above lines indicated significant positive departure from zeros whereas those labeled below represented being smaller than zeros. (**D**) CCH occupancy in [0, 15] ms was correlated with the firing rates of target unit for both subjects (Left: B11, p = 0.01; Right: C05, p = 9e-4). However the correlation showed opposite directions.

Although we have shown that energy-efficient temporal patterns can be generated in M1, it was unclear whether the efficiency was only preserved between direct neurons or performed in a broader range. Therefore, we investigated the excitation level of indirect units (units that were recorded but not used for direct BMI control). We first collected the indirect units exhibiting significant modulation in their firing rates and computed their d-MIs. We found that in the last four sessions when behavioral performance plateaued, the overall d-MI of significant modulated units were either dominated by positive d-MI or counterbalanced (Fig. 5C). The neurons up-modulating their spiking activity did not rule the network. Therefore, we concluded that efficiency was achieved at the population level.

To further explore whether there remained any synergetic evolution between temporal precision and firing rates modulation, we fitted a linear model between the CCH occupancy of [0, 5] ms and firing rates of the target unit, and revealed that those two factors co-evolved for both subjects (Fig. 5D). However, the firing rates of the trigger unit did not evolve with them (Supp Fig. 1).

## Discussion

In this study, we proposed a novel brain-machine interface-based task to investigate the implementation of energy-efficient temporal patterns. We demonstrated that subjects could learn the task and volitionally generate energy-efficient temporal patterns in a reproducible manner in M1, a brain area containing rich neural dynamics to direct motor output. Notably, generating these patterns did not invoke global excitation. Therefore, this novel paradigm can serve as the basis for understanding neural codes under energy constraints and for inspiring energy-saving neuromorphic systems.

Despite the efficiency, another notable disparity between this article and our previous study (Ning et al., 2022) was the temporal order. We can see from Fig. 3 that the distribution of NCS expanded bilaterally which suggested an insensitivity towards the temporal order of spikes. Since neither “decoder” has posed constraints on the neural patterns or dynamics when subjects were approaching the target patterns, we postulated this disparity might arise from either the between efficient and non-efficient temporal patterns or the structural differences between species. Further studies are needed to disentangle these factors.

There was a marked increase in the modulation depth of target neurons shown in our results (Figure 5). However, the two subjects exhibit different evolutionary dynamics of firing rates. Specifically, the mean firing rates increased for Monkey B11 while steadily decreasing for Monkey C05. Despite the disparity in trends, two subjects converged in average firing rates (Fig. 5B) and the coincidence level (Fig. 4A) in the final stage of learning (Fig. 6). This convergence prompted the plausible speculation that the modulation of firing rates and temporality might be inseparable when generating energy-efficient temporal patterns. If they could be modulated independently, then predictably, B11 could rapidly refine its efficient temporal patterns through slight adjustments in temporal precision between direct neurons. The idea of differential involvement of spike synchronization and rate modulation was first illustrated in (Riehle et al., 1997). According to their argument, synchronization occurs as long as there are stimuli or expected to be one. In the language of probability (Pouget et al., 2000; Quiroga and Panzeri, 2009), the posterior probability of finding external or internal presentation of stimulus *s* given a synchronization *r*, denoted as *P*(*s*|*r*), is high. Thus synchronization was said to encode information of the presentation of the stimulus. However, the presence of an external or internal stimulus was a man-made (posterior/manufactured) concept that fell into the fallacy of encodingism (Brette, 2019). It also holds when asserting modulation of firing rates exclusively encoded the external stimulus. inevitably lead to conclude synchronization and rate modulation are dissociable without noticing the fact that this dissociability is already being preconceptually included in the movement parameters (or stimulus parameters in the context of sensory coding). Based on their results, one can also justify that sheer synchronization and synchronization in the presence of rate modulation are indeed two independent modalities. Therefore, synchronization and rate modulation can be dissociable in the context of probability and information theory but may be coupled in the context of implementation due to the biological constraints suggested by our study using the BMI-based novel paradigm.

**Figure 6.**
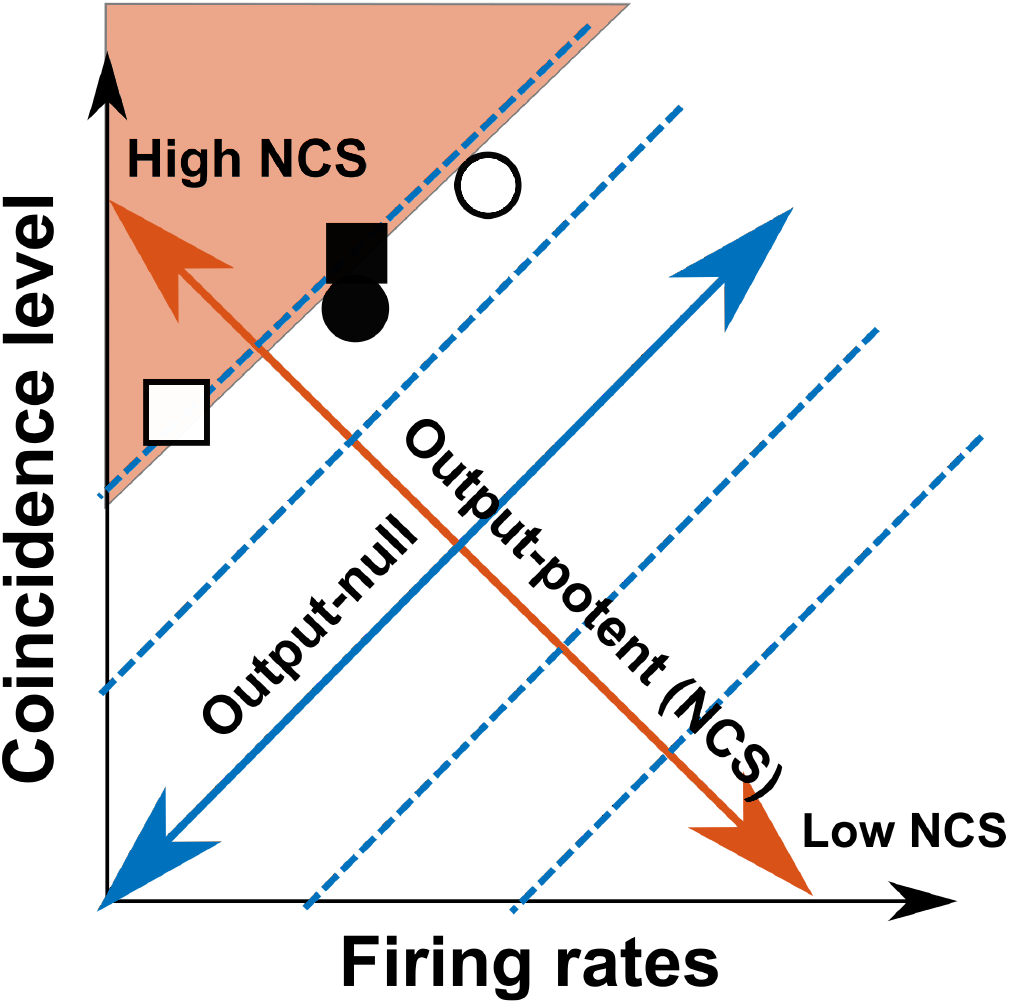
Illustration showing the transition of neural states of the paired direct units featured by their average firing rates and coincidence level. The orange arrow characterized the direction on which the of neural states moving would lead to outcome changes. Orthogonal arrow in blue characterized the direction on which the of neural states moving would not lead to outcome changes. Therefore, any movement in state space could be decomposed by these bases. Squares and circles represent monkey B11 and C05, respectively. The open marks represented the neural states of the first sessions, while the filled marks represented the last sessions.

How did this newly-proposed task paradigm prevail in uncovering the implementation of efficient temporal patterns? User-defined mappings (the “decoder”) of BMI separated the whole neural space into two orthogonal subspaces. Navigating through the “output-potent” space gives rise to immediate changes in the behavioral outcomes while the “output-null” delineating a redundant space has no direct impact. However, several pioneering studies have uncovered organized patterns in redundant space. Often, this neural redundancy manifests pertinent neural functions ((Kaufman et al., 2014; Li et al., 2016; Driscoll et al., 2017)) or circuit constraints in implementing the target neural patterns ((Hennig et al., 2018)). Our study also benefited from this categorically defined redundancy. Here we characterize the neural space through two major features of neural patterns: temporal precision and firing rates, rather than as each dimension by convention, then we obtain the “output-potent” and “output-null” spaces as well (Fig. 6). Further, a broad “reward-null” space was left for subjects to sail across unconstrainedly (the orange area in Fig.6). This area covers all neural patterns that earn rewards indistinctly. Nevertheless, from the results presented in Fig. 5D, the subjects evolved with rather linear trajectories through this “reward-null” space against many theoretically possible routes. Besides, the trajectories converged in the same endpoint in the “reward-null” space regardless of their different initial conditions (Supp Fig. 2). Therefore, this newly-proposed task paradigm is a powerful tool by introducing redundant space. When engaging more initial conditions and their yielded trajectories, we expect to characterize the complete evolution of neural states in learning and explore the underlying mechanism in future works.

Ever since Shannon proposed the renowned information theory (Shannon, 1948), neuroscientists have applied it to estimate the entropy of spike trains (MacKay and McCulloch, 1952; Strong et al., 1998; Rieke et al., 1999; Tang et al., 2014; Pryluk et al., 2019). However, when calculating this entropy, patterns (binary strings) that differed in spikes placement but equaled in the number of spikes were assigned uniform probability. This contrasted to the sparse occupation of pattern space that our results have indicated and thus rendered an overestimation of temporal coding capacity that a pair of neurons could carry. Therefore, we need to refine the estimation of information capacity of spikes with biologically informed priors and revisit what is the arena for temporal coding to manifest its advantages. For example, their relative spike timing allowed multiplexing of stimulus features (Hopfield, 1995; Gawne et al., 1996; Brody and Hopfield, 2003). Besides, though the limited capacity to carry information was suggested for neuron pairs, the capacity might be preserved in the neural population (Gollisch and Meister, 2008).

Unlike previous research that characterized how efficient temporal codes correlated with overtrained behaviors or reproducible natural behaviors, BMI paves the way for probing neural dynamics throughout learning since the causal relationship between neurons and outcomes is overt. Therefore, the theoretical moment-by-moment or trial-to-trial errors could be explicitly inferred, which allowed quantitative tests of learning models or credit assignment models.

## Methods

### Surgery and Data Acquisition

All surgical and experimental procedures conformed to the Guide for The Care and Use of Laboratory Animals (China Ministry of Health) and were approved by the Animal Care Committee of Zhejiang University, China. The surgical procedure was described in detail in (Zhang et al., 2012). In brief, the 96-channel microelectrode arrays (Blackrock Neurotech) were chronically implanted in the primary motor cortex of two male rhesus monkeys (Macaca mulatta) (Monkey B11, Monkey C05). The monkeys took roughly a week to recover from the surgery, after which the neural signals were recorded through the Cerebus multichannel data acquisition system (Blackrock Neurotech) at a sample rate of 30 kHz. Spike activities were detected by thresholding (root mean square multiplier, B11: ×5.5; C05: ×7). On the first session of learning, after online manual sorting, we assigned two isolated units with the highest signal-to-ratio (SNR) to be the trigger unit and the target unit, respectively. The online sorting templates for these two units were unchanged throughout the learning and validated by offline sorting (Plexon, Inc.). Since these two selected units were directly responsible for the outcome, they were labeled as “direct neurons”. The remaining units were “indirect neurons”, which were sorted offline for the analysis in this article.

### Behavioral task

In this task, the spikes of the trigger unit and the target unit in a window of 300 ms were used to compute a conditioning variable: normalized coincidence score (NCS). This window was slid every 150 ms in real-time. NCS was then fed back to the subject by audio and vision. The subjects would receive the water reward if they up-modulated the NCS to a threshold in 15 s. Otherwise, the trial would be aborted, and the subjects should wait another 4 s for the next trial (Fig. 1B). Thresholds of NCS for obtaining rseward were set to be the 99^th^ percentile of the NCS distribution estimated from the baseline data at the first session (sampled every 150 ms for 5 minutes). These thresholds (B11: 0.36; C05: 0.32) were fixed across sessions.

We defined the conditioning variable NCS which measured the spike coincidence level as:

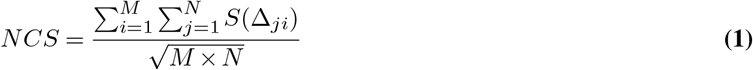

where *M* and *N* were the total numbers of spikes from trigger unit and target unit in one data bin, *S*(Δ*_ji_*) was the score function dependent on the lag *Δ_ji_* between the time of the *i^th^* spike of the trigger unit and the *j^th^* spike of the target unit. Specifically, the score function took the similar exponential form as how synaptic efficacy changes under spike timing-dependent plasticity (STDP) (Dan and Poo, 2004), where the trigger leading target resulted in a postive score and vice versa:

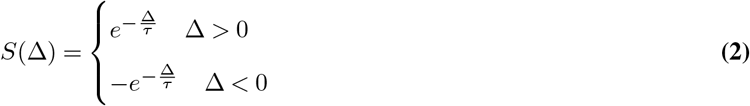

*τ* was set to 15 ms to fit the critical Window observed in STDP experiments. Therefore the value of NCS should be in the range from −1 to 1.

We mapped the NCS into the frequency of an audio cursor ranging from 1 Hz to 24 kHz in quarter-octave increment. Also, we adapted a similar visual feedback environment since all subjects had once been trained on the center-out tasks. NCS in [-0.5, 0.5] was mapped to the vertical position of a blue ball on the screen (Fig.1A), with a yellow ball cue for the target.

Subjects were water-constrained and learned tasks on consecutive days, each containing two sessions in the morning and afternoon.

### Data Analysis

We used two metrics: normalized success rate and trial duration, to evaluate the task performance. The raw success rate *SR_raw_* was defined as the ratio between the number of successful trials and the total number of trials in one session. Assuming that the random process *NCS*(*t*) under chance level was stationary, then the chance level of the raw success rate *SR_CL_* was the average probability of exceeding the threshold in 15 s, denoted as *P_trial_*. We could estimate the probability density function *f*(*NCS*) by uniformly sampling NCS in the Baseline. The corresponding percentile of the given threshold was denoted as *P_thr_* which characterized the probability of failing to exceed the threshold in one bin (300 ms). Therefore, concerning that failing one trial follows a binomial distribution with 50 independent experiments, we have:

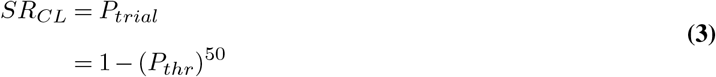

To counter the variation of chance level, we normalized *SR_raw_* by *SR_CL_* to render a normalized success rate *SR_norm_*:

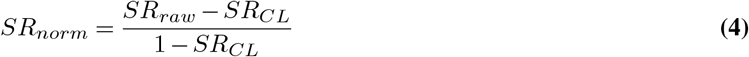

Trial duration was defined as the time from the start of a trial to the reward delivery.

We used a jitter-corrected cross-correlation histogram (CCH) to portray the temporal relationship between the trigger and target units on a fine time scale (Amarasingham et al., 2012; Vizuete et al., 2012). We randomly jittered the emitting time for each spike from the target unit to render the surrogated CCH (sCCH). This resampling procedure was run 1000 times. The jittering variation was dependent on the temporal resolution of the CCH. For example, if the bin of CCH was 15 ms as shown in Fig. 4A, then the standard deviation of random jittering (“jitter window”) should also be set as 15 ms. We subtracted this sCCH from the crude CCH to obtain the corrected CCH. In this way, we guaranteed that structured firing patterns or variation in timescales comparable to the temporal resolution of CCH would be removed. The tail probability of sCCH was used to construct the acceptance bands. The error bar in the jitter-corrected CCH outlined the standard deviation of sCCH.

The directional modulation index (d-MI) was used to characterize the modulation depth for single neurons in the task block grounded on firing in the baseline block *FR_baseline_*:

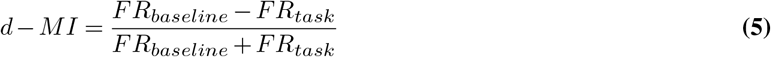

Overshooting NCS were those who passed the threhold for rewarding, as well as the last computed NCS in a successful trial. To counter the disparity of threshold values in two subjects, we subtracted the difference in thresholds (0.04) from all the overshooting NCS of B11 to render the corrected overshooting NCS.

## Data and Code Availability

Data and codes will be made available upon reasonable request at the time of publication.

## Author Contributions

Y.N. and S.Z. conceived and designed the study. Y.N. and G.W performed the animal experiment. Y.N. and T.L. performed the analysis. Y.N. wrote the manuscript with input from T.L. and S.Z.

## Supplementary Information

**Figure S1.**
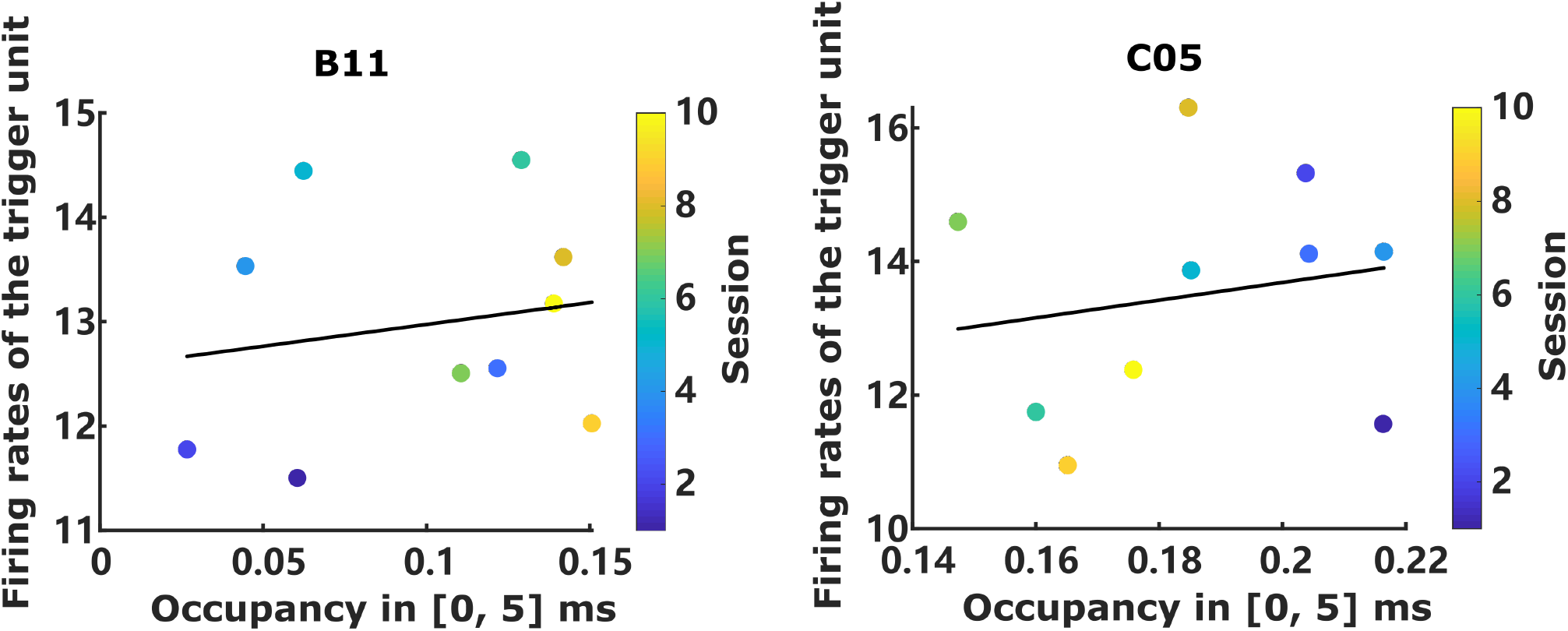
The firing rates of trigger unit was not correlated to the temporal precision across sessions. *Left*: p = 0.62, *right*: p = 0.61

**Figure S2.**
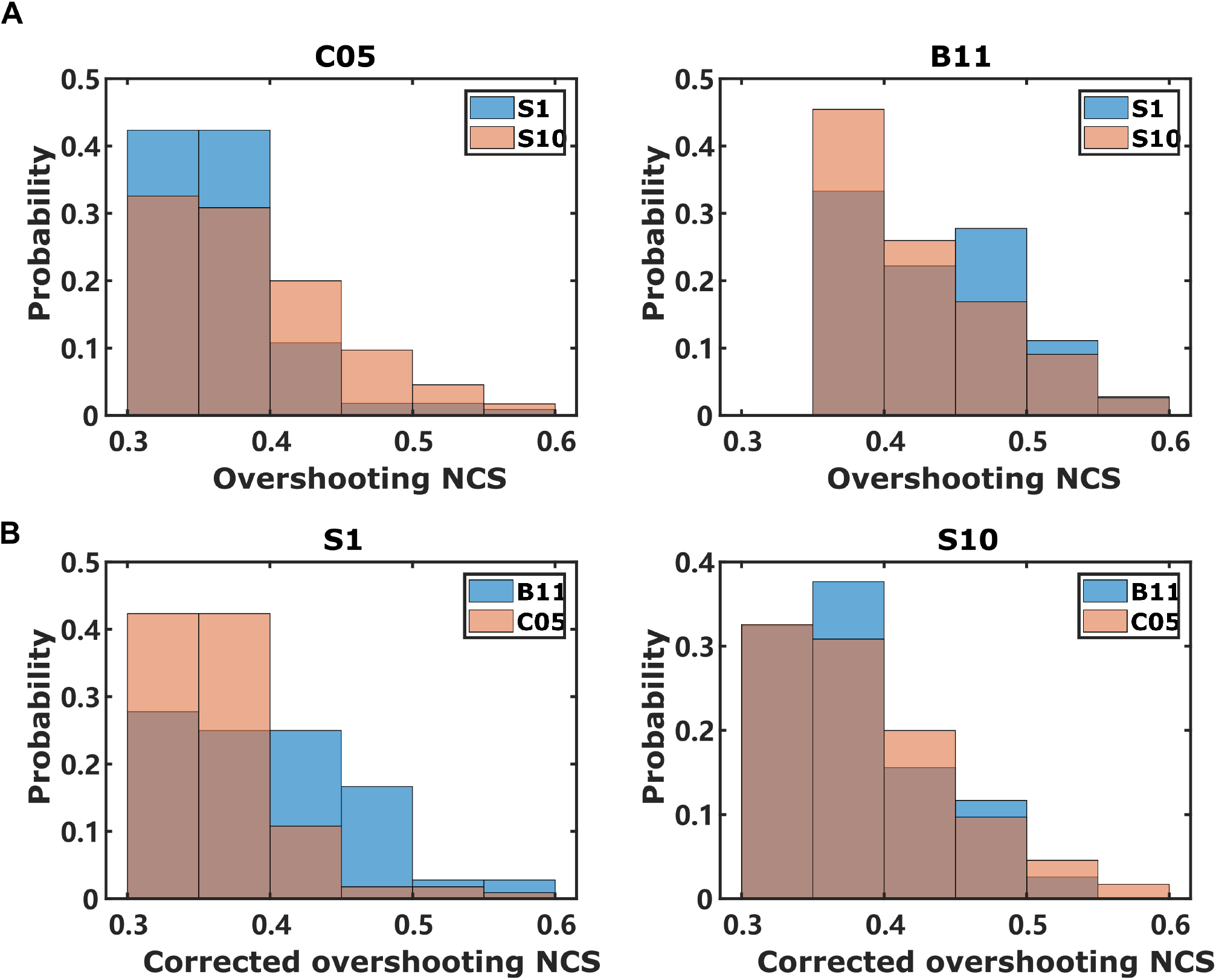
Distribution of overshooting NCS before and after learning. (**A**) Histograms of overshooting NCS from Session 1 (*blue*) and Session 10 (*Orange*). (**B**) Histograms of overshooting NCS after correcting from Session 1 (left) and Session 10 (*right*). Subjects are color-coded.

